# Spatially resolved transcriptomics identifies intercellular signaling post-ischemic stroke that controls neural stem cell proliferation

**DOI:** 10.1101/2025.09.04.674219

**Authors:** He Huang, Emerson Daniele, Wing Chung Jessie Lam, Teodora Tockovska, Daniela Lozano Casasbuenas, Hathairat Chanphao, Bebhinn Treanor, Maryam Faiz, Scott A. Yuzwa

## Abstract

Stroke is the second leading cause of death and disability worldwide. Ischemic stroke mobilizes adult neural stem cells (NSCs) out of the quiescent state. The multifaceted responses of endogenous NSCs to ischemic stroke involve proliferation, migration, and differentiation. Hence one strategy which could be leveraged for recovery after ischemic stroke is the intrinsic mechanism of endogenous NSC mobilization. However, the survival rate of recruited endogenous NSCs is low. Moreover, the intercellular signals that activate NSCs after ischemic stroke are poorly understood. We hypothesized that after stroke, cells located in the cerebral lesion send signals to the NSC niche to initiate the regenerative response. To test this hypothesis, we used CellChat to computationally infer the cell-cell communication between the ischemic infarct region and ventricular-subventricular zone (V-SVZ) NSC niche from spatial gene expression profiles. We identified ligand-receptor pairs and signaling pathways involved in the signal transduction events at 2, 10, and 21 days after stroke. Out of several candidate genes of interest we identified, here we reported the regulatory function of galectin-9 on the proliferation of NSCs. Our present work portrays galectin-9 as a checkpoint signaling molecule that guards the responses of NSCs under physiological conditions and potentially during the recovery phase post-ischemic stroke. We suggest that TIM-3 mediates the inhibitory effect of galectin-9 on NSC proliferation and propose a working hypothesis that the stroke-induced proinflammatory factors stimulate the Toll-like receptor 4 (TLR4) on ependymal cells and result in the increased secretion of galectin-9, which in turn modulates neighboring NSCs. Our study paves the way for potential therapeutic approaches which leverage the TLR4 and galectin-9/TIM3 signaling pathways.

## INTRODUCTION

Globally, stroke is the second most prevalent cause of death and disability ^1^. Ischemic stroke accounts for 85% of the total cases with the remaining 15% being hemorrhagic ^1^, both of which can be further divided into various subtypes according to different classification systems based on the etiologies and phenotypes ^2^. Ischemic stroke has been studied extensively using numerous animal models, including middle cerebral artery occlusion (MCAO), craniotomy, photothrombosis, endothelin-1 (ET-1), and embolic stroke models, the details of and pros and cons of which have been thoroughly reviewed ^3^.

Ischemic stroke is a brain injury that triggers the reactions of neural stem cells (NSCs) ^4^. NSCs are characterized by their capability of self-renewal and to produce differentiated neural cells via asymmetric cell division ^5,6^. In the adult mammalian brain, NSCs exist in specific niches of the ventricular-subventricular zone (V-SVZ) of the lateral ventricle wall and the subgranular zone (SGZ) of the hippocampal dentate gyrus ^7^. The multifaceted responses of endogenous NSCs to ischemic stroke ^8^ involve proliferation ^9^, migration ^9^, differentiation ^10^, and synaptogenesis ^11^. Previous studies have shown that in adult rodent brain, following ischemic insults, neurogenesis occurs from the neural stem and/or progenitor cells in the subventricular zone, or local cortical progenitors, or striatal astrocytes ^12^. However, around 80% of the new neurons formed in the injured striatum die between 2 and 6 weeks post-ischemia ^9^. Post-mortem studies on adult human brain of patients after ischemic stroke revealed increased numbers of proliferating cells in ipsilateral SVZ in comparison with the contralateral SVZ ^13,14^ and uninjured control ^13^. Although stem cell therapies have shown promising potential for treating ischemic stroke ^15^, some well-recognized risks such as tumorigenesis have been documented ^16^. Intuitively, another strategy for promoting recovery after ischemic stroke is to take advantage of the intrinsic mechanisms of endogenous NSC mobilization. However, the intercellular signals which activate NSCs after brain injury are still incompletely understood.

Intercellular signaling can be inferred by spatial transcriptomics ^17,18^ that enables high-throughput quantification of gene expression *in situ*, and therefore characterization of cell-cell communication via defined ligand-receptor (L-R) pairs ^19^. Many tools and resources have been developed and curated for deciphering cell-cell communication, which were systematically reviewed and compared recently ^20^. In spite of differences in strategies adopted among these methods, all infer cell-cell communication from gene expression, predominantly focusing on L-R pairs ^21^. We hypothesized that after stroke, the cells located in the cerebral infarct secrete and send signals to the NSC niche to initiate the regenerative response.

Here we used CellChat ^22^ to computationally infer signaling between the ischemic infarct region and V-SVZ NSC niche. We identified L-R pairs and signaling pathways potentially involved in the signal transduction events 2, 10, and 21 days after ischemic stroke. Many of the L-R pairs have previously been characterized as regulators of NSC proliferation, migration, differentiation, and synaptogenesis. After screening several candidate genes of interest with limited known roles in the NSC niche, our current study presents the first evidence, to our knowledge, of the regulatory function of galectin-9 on the proliferation of NSCs from both *in vitro* and *in vivo* assays as well as the galectin-9 knockout (*Lgals9*^−/−^) mouse. Galectin-9 is a β-galactoside binding lectin ^23^ that has not only been extensively studied as an immune response modulator ^24–26^ but also associated with the post-stroke recovery ^27–31^. Our present work depicts galectin-9 as a checkpoint signaling molecule that under physiological conditions controls NSC proliferation and this physiological pathway is likely leveraged in the later recovery phase post-ischemic stroke. We further investigate the potential receptors for galectin-9 identified by our spatial transcriptomics data and suggest that TIM-3 mediates the inhibitory effect of galectin-9 on NSC proliferation. We therefore propose a working hypothesis that upon the stimulation of the Toll-like receptor 4 (TLR4) on ependymal cells in the V-SVZ of the adult mouse brain, the activated ependymal cells, which physiologically produce galectin-9 to control NSC proliferation, increasingly produce galectin-9 after ischemic stroke, which in turn modulates the NSC niche.

## RESULTS

### Intercellular communication inference from spatial transcriptomics

To characterize signal transduction between the infarct of ischemic stroke and the NSC niche (Figure 1A), we started by computationally inferring intercellular communication from spatial transcriptomics data. Two 10x Visium platform datasets were analyzed, each of which had three time points – acute (day-2), sub-acute (day-10), and chronic (day-21) post-ischemic stroke, along with a control sham injury (day-2 sham), as we described previously ^32^. The stroke infarct regions and the saline injection site of sham control were manually delineated with the 10x Genomics Loupe Browser. To identify NSCs, the barcoded spots in the V-SVZ were hand-selected with the 10x Genomics Loupe Browser. The sum of the normalized counts of the NSC marker genes – *Sox2*, *Nes*, *Gfap*, *Prom1*, and *Fabp7* was calculated and then ranked across all spots. The identity of NSC was assigned to spots in the first quartile of the ranks. We used CellChat ^22^ to infer cell-cell communication. To get an overview of the changes of intercellular signaling during the time course post-ischemic stroke in our animal model, we began with comparing the total number of interactions and interaction strength for samples of each time point and the sham control. The total numbers of intercellular interactions and interaction strength were increased in day-2 and further increased in day-10 samples compared with sham but then decreased by day-21 (Figure 1B). This pattern agreed with the previous characterization of endothelin-1 (ET1) induced stroke in rodents where the most neurological deficits were observed in one week post ischemic-stroke followed by a significant recovery phase ^33,34^. To reveal the signaling molecules and pathways that play important roles in mobilizing NSCs located in the V-SVZ niche, we focused on the interactions (ligand-receptor pairs) from the stroke infarct site to NSCs (Figure 1C). The enhancement of cell-cell communication post-ischemic stroke was visually striking using chord diagrams to represent individual ligand-receptor (L-R) pairs.

**Figure 1.**
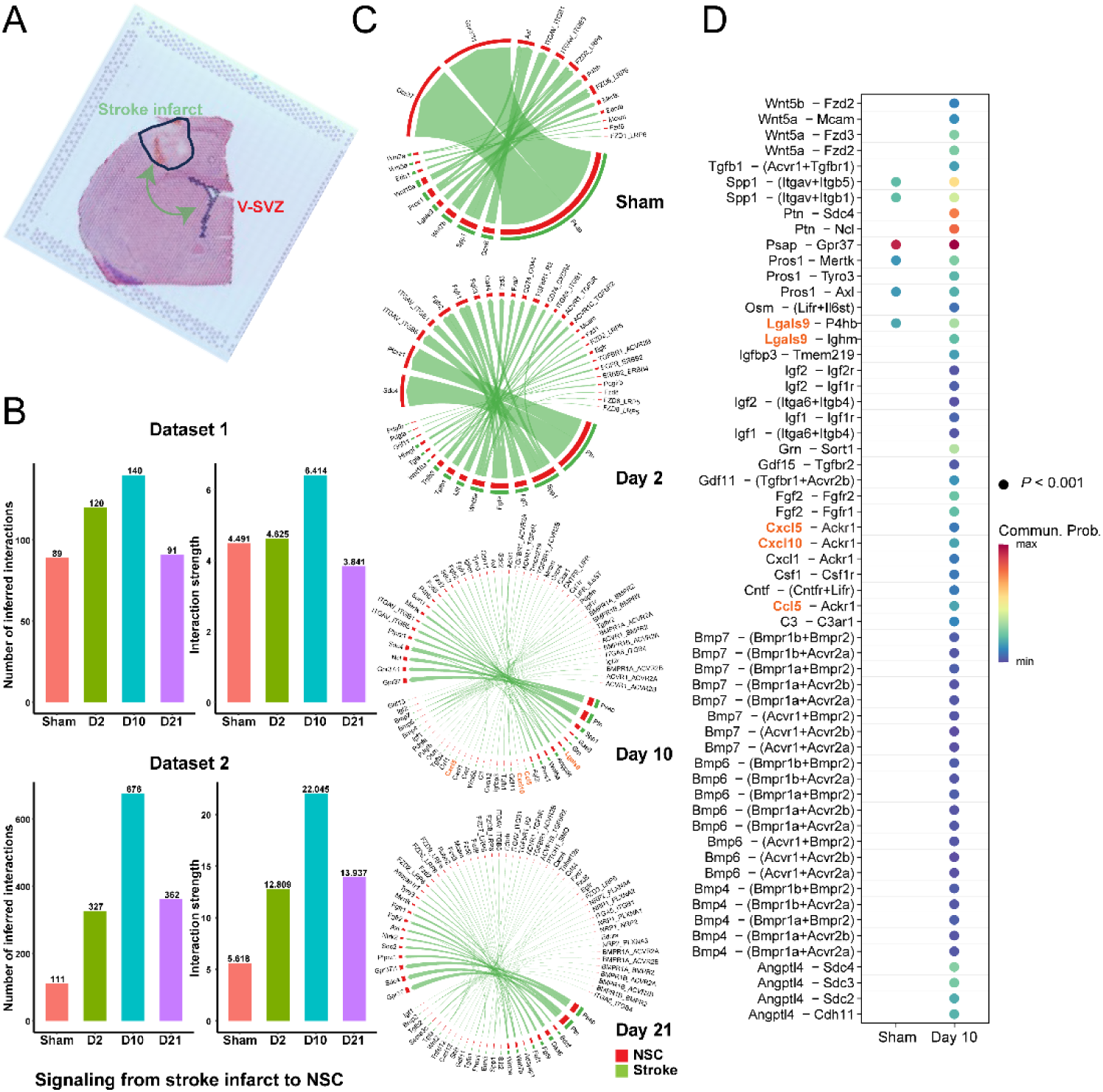
Inference of cell-cell communication from spatial transcriptomics data. (A) Hematoxylin and eosin stained section on a Visium spatial gene expression capture area ^32^. The cell-cell communication between cortical ischemic lesion produced by endothelin-1 (ET-1) injection and the V-SVZ NSC niche was inferred by CellChat. (B) The comparison of the total number of interactions and interaction strength among sham control, day-2, day-10, and day-21 post-ischemic stroke. (C) The ligand-receptor pairs and intercellular signaling from the stroke infarct site to the V-SVZ NSC niche inferred from spatial transcriptomics data in sham control and three time points post-ischemic stroke. (D) The upregulated intercellular signaling ligand-receptor pairs from stroke infarct to V-SVZ NSCs in 10 days post-ischemic stroke when compared to sham control by comparing the communication probabilities (*P* < 0.001).

Next, we conducted a pairwise comparison of the up– and down-regulated signaling L-R pairs based on the communication probabilities (Table S1). Compared with sham, the numbers of upregulated L-R pairs were 128, 387, and 199 in day-2, day-10, and day-21 sample, respectively. Moreover, 342 L-R pairs were upregulated in day-10 in comparison with day-2 sample, whereas 385 L-R pairs were down-regulated in day-21 compared to day-10. These numbers clearly suggested that the signal transduction from stroke infarct to NSC rose in the acute phase after stroke (day-2), peaked around sub-acute phase (day-10), and started to decline in the recovery/chronic phase (day-21). To pinpoint the critical regulators of NSCs post-stroke, we focused on the increased signaling from stroke lesion to NSCs in 10 days post-ischemic stroke when compared to sham (Figure 1D). Particularly, the L-R pairs upregulated in day-10 compared with both sham and day-2 but downregulated compared with day-21 were considered for further analyses.

Many of the intercellular signaling events identified computationally have been previously regarded as regulators of NSC proliferation, migration, differentiation, and synaptogenesis, such as *Psap* ^35^, *Ptn* ^36^, and *Grn* ^37^ which validated our bioinformatic analysis. To expand the knowledge of cell signaling from stroke infarct to NSCs, we explored four ligands inferred from our computational analysis but not previously well characterized in this context, including *Lgals9* (galectin-9), *Ccl5* (C-C motif chemokine ligand 5), also known as RANTES (Regulated upon Activation Normal T cell Expressed and Secreted), *Cxcl5* (C-X-C motif chemokine ligand 5), also known as ENA78 (Epithelial cell-derived Neutrophil-Activating peptide-78), or LIX (LPS-Induced CXC chemokine), or GCP-2 (Granulocyte Chemotactic Protein-2), or AMCF-II (Alveolar Macrophage Chemotactic Factor-II), and *Cxcl10* (C-X-C motif chemokine ligand 10), also known as IP-10 (Interferon γ-induced Protein 10 kDa). The spatial expression patterns of these four ligands demonstrated the same trend as the overall numbers and strength of intercellular communication, with salient increase in 10 days post-ischemic stroke and focal expression at the stroke infarct site in day-10 sample. (Figure 2A). To identify the types of cells that generated these signaling molecules, we analyzed an existing published single-cell RNA-sequencing dataset where the cell types were annotated by known cell type markers (Figure 2B) ^38^. As a well-known immunomodulator ^24–26,39^, in agreement with other data sources ^40–42^, galectin-9 was primarily expressed in microglia (Figure 2B). Noticeably, galectin-9 expression also presented in ependymal cells. CCL5 was mainly expressed in microglia. CXCL10, another chemokine, was found in both microglia and ependymal cells. CXCL5 was highlighted in the oligodendrocyte lineage.

**Figure 2.**
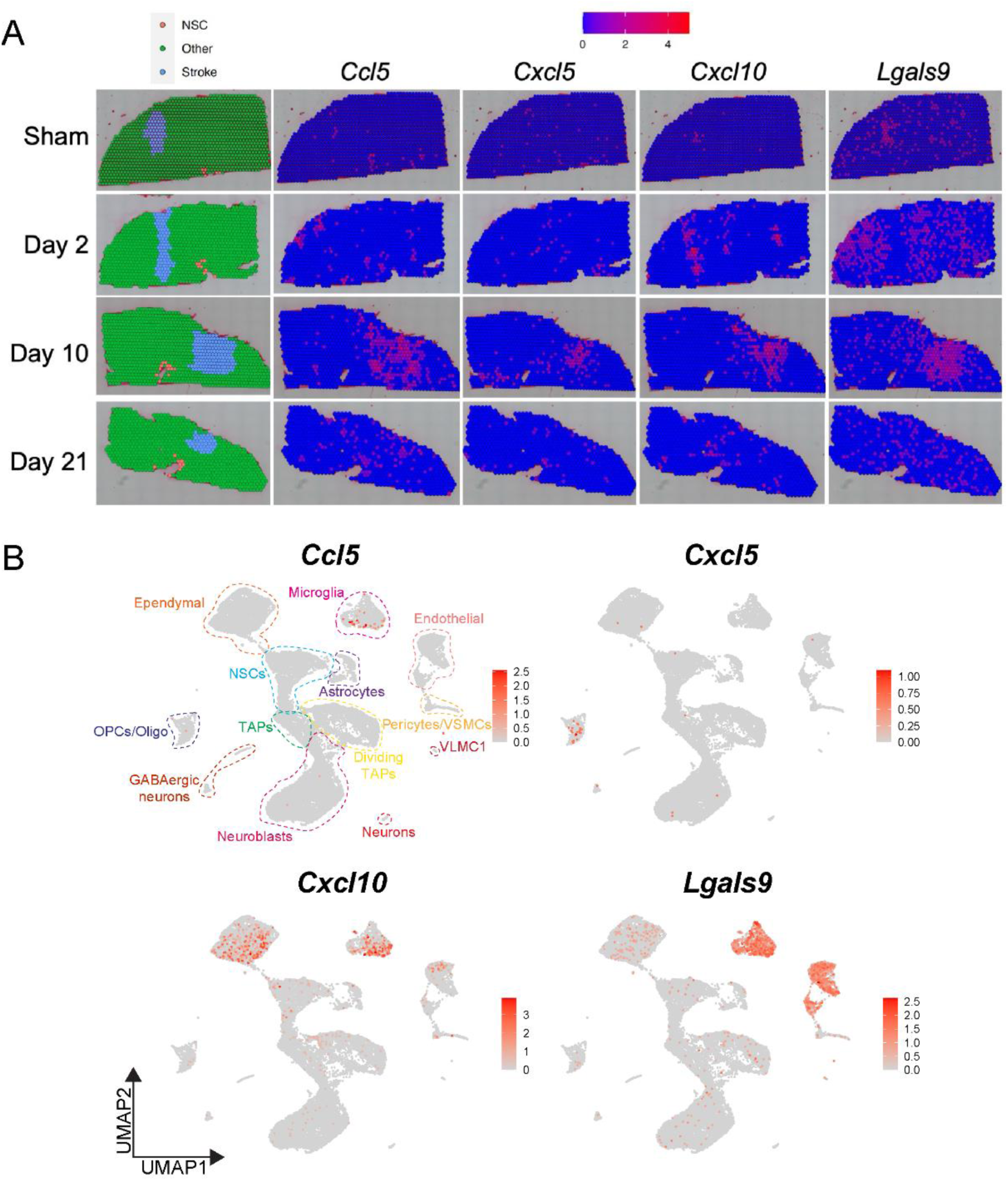
The expression pattern of selected genes of interest. (A) The expression patterns of selected genes of interest from 10x Visium spatial transcriptomics data from brain sections of sham control, day-2, day-10, and day-21 post-ischemic stroke. (B) The expression patterns of select genes of interest on a UMAP plot of V-SVZ whole-cell single-cell analysis from Cebrian-Silla *et al*. ^38^. The cell types were identified by marker genes ^38^.

### Intercellular communication inference identifies potential regulators of NSC proliferation

One of the possible functions of these signaling molecules elicited post-ischemic stroke could be to enhance NSC proliferation in the V-SVZ niche. To ask whether these ligands might be unrecognized modulators of NSC proliferation post-ischemic stroke, we first surveyed the impacts of the four ligands identified computationally *in vitro* using neurospheres (Figure 3). We evaluated the effects of four ligands on the proliferation of the secondary spheres, including galectin-9, CCL5, CXCL5, and CXCL10. NSC proliferation was evaluated indirectly using the resazurin reduction assay and quantified by the fluorescence intensity of resorufin ^43^. We observed significant impacts of the ligands on NSC proliferation (Friedman test, χ^2^ = 14.63, *P* = 0.0055, *N* = 7). Galectin-9 decreased NSC proliferation by nearly 2-fold compared to the control (Dunn’s multiple comparisons test, *Adj. P* = 0.0029), whereas the other three ligands tested had no significant effects on NSC proliferation (Figure 3B). To confirm the inhibitory effects of galectin-9 on NSC proliferation, we quantified the sizes of neurospheres. The diameters of neurospheres cultured with galectin-9 showed approximately 20% reduction compared with those of control (paired *t*-test, *t*_(9)_ = 2.663, *P* = 0.0259) (Figure 3C), which when converted to volume reflects a ∼2-fold reduction of the volume of neurospheres (Figure 3A), which compared favorably to the resazurin reduction assay data (Figure 3B). To further examine how galectin-9 influenced the stemness of NSC, we quantified the counts of these secondary neurospheres. Galectin-9 trended towards slightly increased numbers of neurospheres (paired *t*-test, *t*_(9)_ = 1.824, *P* = 0.1014) compared with the control (Figure 3D) but this did not reach significance, suggesting relatively little impact on the stemness of NSCs. Collectively, our *in vitro* experiments demonstrated that galectin-9 inhibited NSC proliferation but preserved NSC stemness.

**Figure 3.**
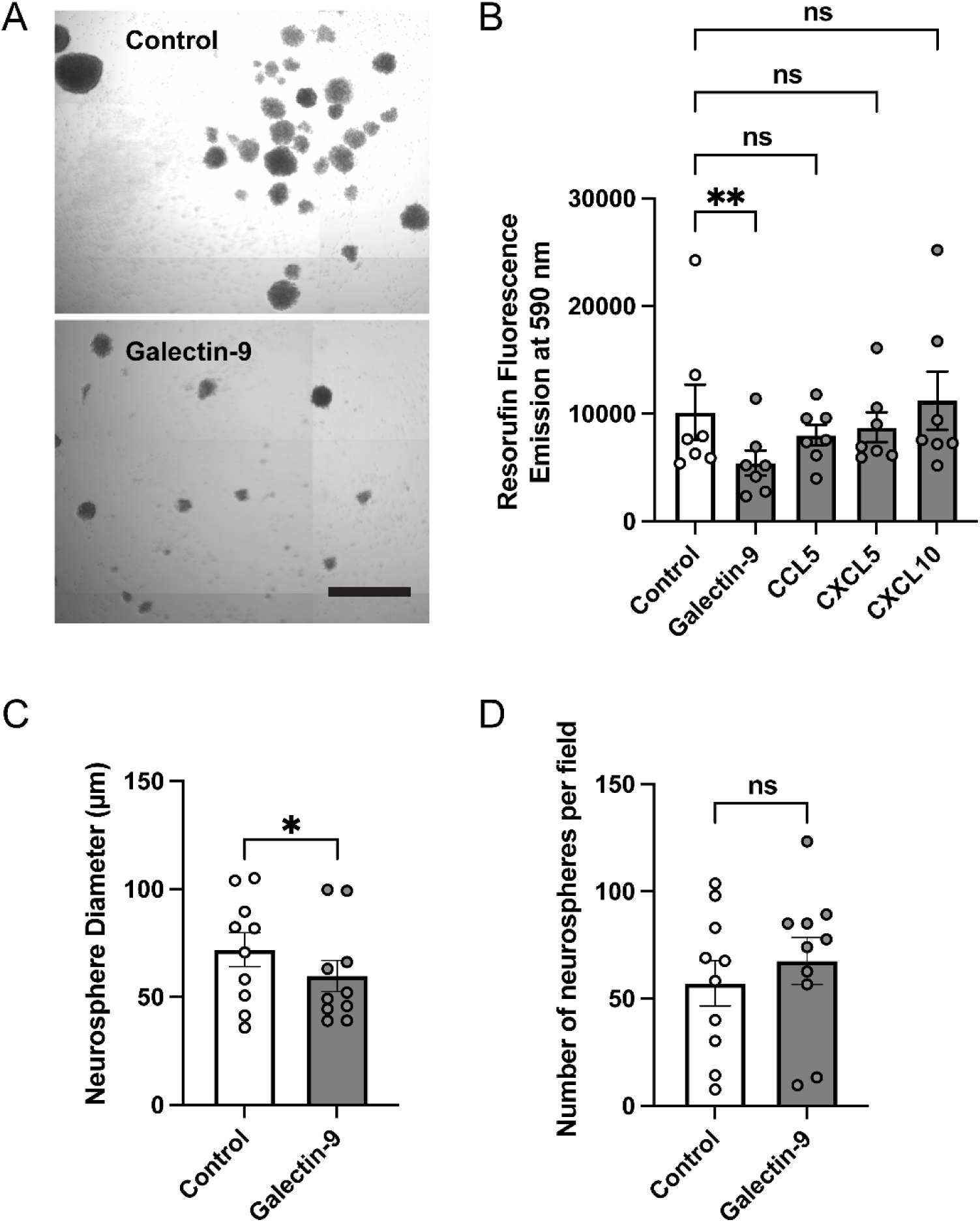
Galectin-9 inhibits NSC proliferation *in vitro*. (A) Comparison of neurospheres cultured with Galectin-9 (2 μg/ml) and control. The scale bar represents 50 μm. (B) The effects of selected signaling molecules on NSC proliferation, which was evaluated by resazurin reduction assay ^43^ and quantified by the fluorescence intensity of resorufin (*N* = 7 per group). (C) Comparison of the diameters of neurospheres cultured with galectin-9 (2 μg/ml) and control (*N* = 10 per group). (D) Comparison of the numbers of neurospheres cultured with galectin-9 (2 μg/ml) and control per microscopic field of the well in a 96-well plate (*N* = 10 per group).

Another potential role of the signaling molecules from the stroke infarct site to the NSC is to induce NSC migration to the injured area as previously demonstrated ^9^. Given the results of our computational inference identified many chemokines, we next evaluated the chemoattractant property for NSCs of these four ligands. We performed quantitative analysis of migration by Boyden chamber assay ^44^. 1 × 10^5^ cells in single-cell suspension disassociated from secondary neurospheres were seeded into each transwell. The cells migrated to the side facing the medium containing galectin-9 (2 μg/ml), CCL5, CXCL5, and CXCL10 (200 ng/ml), along with control, were counted, and expressed as cell numbers per square millimeter (mm^2^). We did not observe significant chemotactic effects of these four ligands on NSCs (Friedman test, χ^2^ = 2.841, *P* = 0.585, *N* = 7) (Figure S1), although CXCL10 mobilized slightly more NSCs than control medium but this did not reach significance. After characterizing the effects of these signaling molecules *in vitro*, our subsequent *in vivo* studies focused on the negative regulatory effects of galectin-9 on NSC proliferation.

### Characterization of the origins and targets of galectin-9 in mouse brain

To further illustrate the mechanism underpinning the inhibitory effect of galectin-9 on NSC proliferation *in vivo*, we investigated the sources and receptors of galectin-9, with a focus on the V-SVZ in the adult mouse brain. Based on the expression of galectin-9 in ependymal cells from single-cell data analysis (Figure 2B), we acquired direct evidence by immunostaining. Our data demonstrated that galectin-9 was detected in FOXJ1 positive ependymal cells along the lateral wall of the ventricles (Figure 4B, arrows). Galectin-9 produced by ependymal cells in the V-SVZ could cause immediate impact on NSCs because of their proximity in this niche. Many receptors of galectin-9 have been previously identified, including for example, TIM-3 (*Havcr2*), TLR4, PD-1, and CD44 ^24^. The bioinformatics inference by CellChat from our spatial transcriptomics data indicated CD44, CD45, TIM-3, IGHM, BCR and P4HB as the receptors for galectin-9 in the signaling pathway activated post-ischemic stroke (Table S1, Figure 1C and 1D). From the analysis of single-cell data, besides microglia, both TIM-3 and TLR4 were expressed in the cell populations in the V-SVZ, including NSCs, transit-amplifying progenitors (TAPs), dividing TAPs, and neuroblasts (Figure 4A). However, TLR4 showed particularly high expression level in ependymal cells, signifying the important roles of ependymal cells in response to injury, infection, and inflammation in the brain, especially given their location lining the ventricles ^45^. Our immunohistochemical data confirmed the expression of TIM-3 in NSCs (Figure 4C, arrows, TIM-3+SOX2+GFAP+ cells) and the expression of TLR4 in both NSCs (Figure 4D, arrows, TLR4+SOX2+GFAP+ cells) and ependymal cells (Figure 4D, arrowheads, TLR4+SOX2+GFAP-cells).

**Figure 4.**
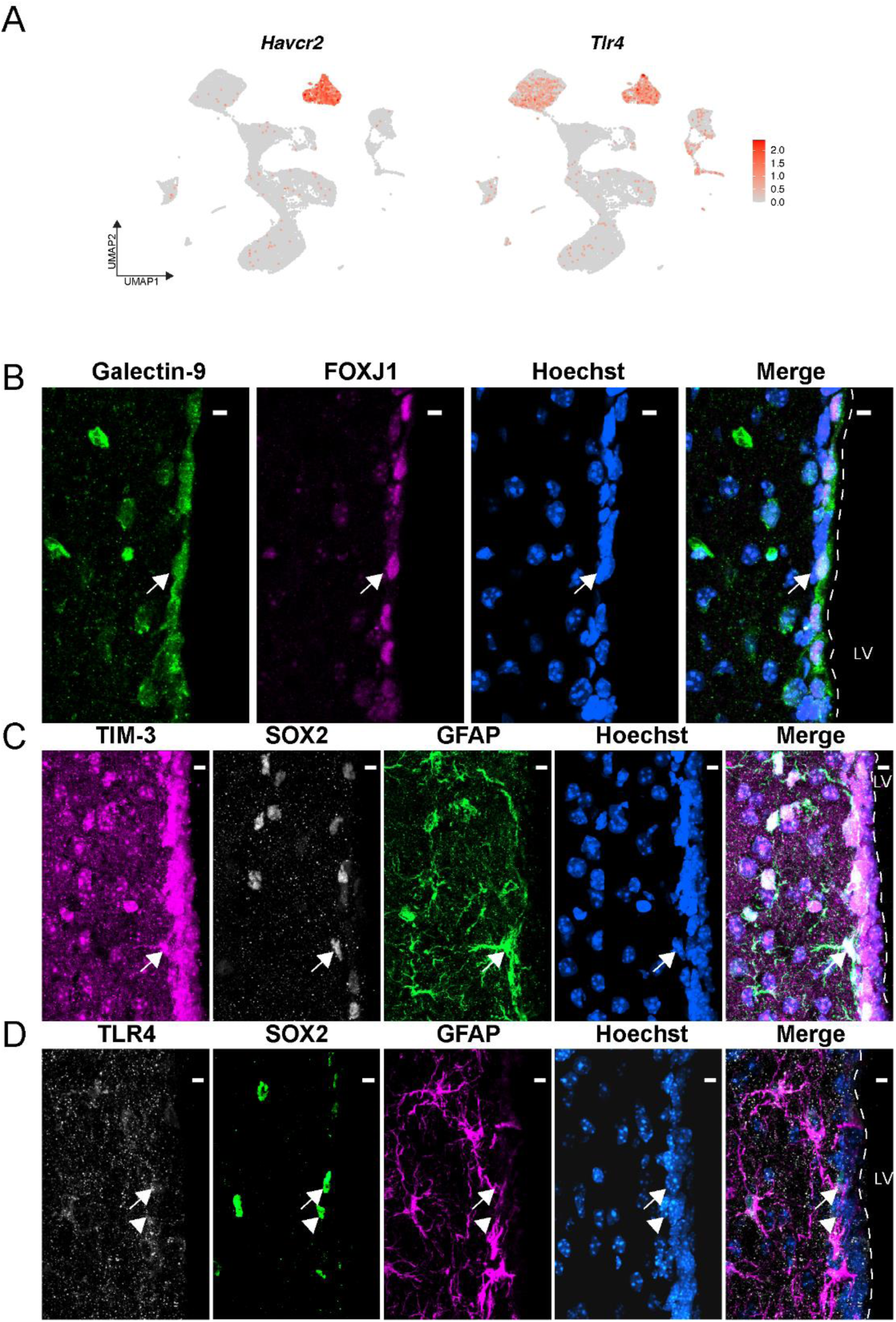
Cell type expression of the galectin-9 and known receptors in the brain. (A) The expression patterns of *Havcr2* (TIM-3) and *Tlr4* on a UMAP plot of V-SVZ whole-cell single-cell analysis ^38^. (B) Immunostaining of the lateral wall of 8-week-old C57BL/6 mouse brains for galectin-9, FOXJ1, a marker gene for ependymal cells, and merged with Hoechst to label nuclei. Arrows indicate the colocalization of galactic-9 and FOXJ1. (C) Immunostaining of the lateral wall of 8-week-old C57BL/6 mouse brains for TIM-3, SOX2, GFAP, and merged with Hoechst to label nuclei. SOX2+GFAP+ cells allow identification of NSCs. Arrows indicate the representative NSC that expresses TIM-3. (D) Immunostaining of the lateral wall of 8-week-old C57BL/6 mouse brains for TLR4, SOX2, GFAP, and merged with Hoechst to label nuclei. SOX2+GFAP+ cells allow identification of NSCs. SOX2+GFAP-cells allow identification of ependymal cells. Arrows indicate the representative NSC that expresses TLR4. Arrowheads indicate the representative ependymal cell that expresses TLR4. Scale bars represent 10 μm. Dotted lines delineate the lateral wall of the lateral ventricle (LV).

### Galectin-9 inhibits NSC proliferation *in vivo*

We further extended our *in vitro* observation of the negative impact of galectin-9 on NSC proliferation via intracerebroventricular (ICV) injection. Galectin-9 or PBS was injected into the lateral ventricle in the right hemisphere of 8-week-old C57BL/6 mouse brains. NSCs were identified by immunohistochemistry as SOX2+ and GFAP+ cells in the V-SVZ region. Ki-67 immunostaining was used as a cell proliferation marker (Figure 5A). The percentage of proliferating NSCs were quantified in two independent cohorts of mice (Figure 5B). For the first cohort, we collected the brains 1 day and 5 days after ICV injection. The percentage of proliferating NSCs was significantly reduced in the mouse brains injected with galectin-9 in 1-day post-ICV compared with PBS injection (unpaired *t*-test, *t*_(7)_ = 2.678, *P* = 0.032). This inhibitory effect remained until 5 days post-ICV injection, but the difference was no longer significant (unpaired *t*-test, *t*_(8)_ = 1.542, *P* = 0.16), possibly due to the elimination of galectin-9 from the lateral ventricle after injection. We repeated the ICV experiment for a second cohort and collected the mouse brains 1-day after injection. The results faithfully recapitulated the significant reduction of the percentage of proliferating NSCs in brains injected with galectin-9 compared to the control (unpaired *t*-test, *t*_(8)_ = 2.519, *P* = 0.036). Our data demonstrated that galectin-9 inhibited NSC proliferation *in vivo*, echoing our *in vitro* evidence obtained from the neurosphere assay (Figure 3).

**Figure 5.**
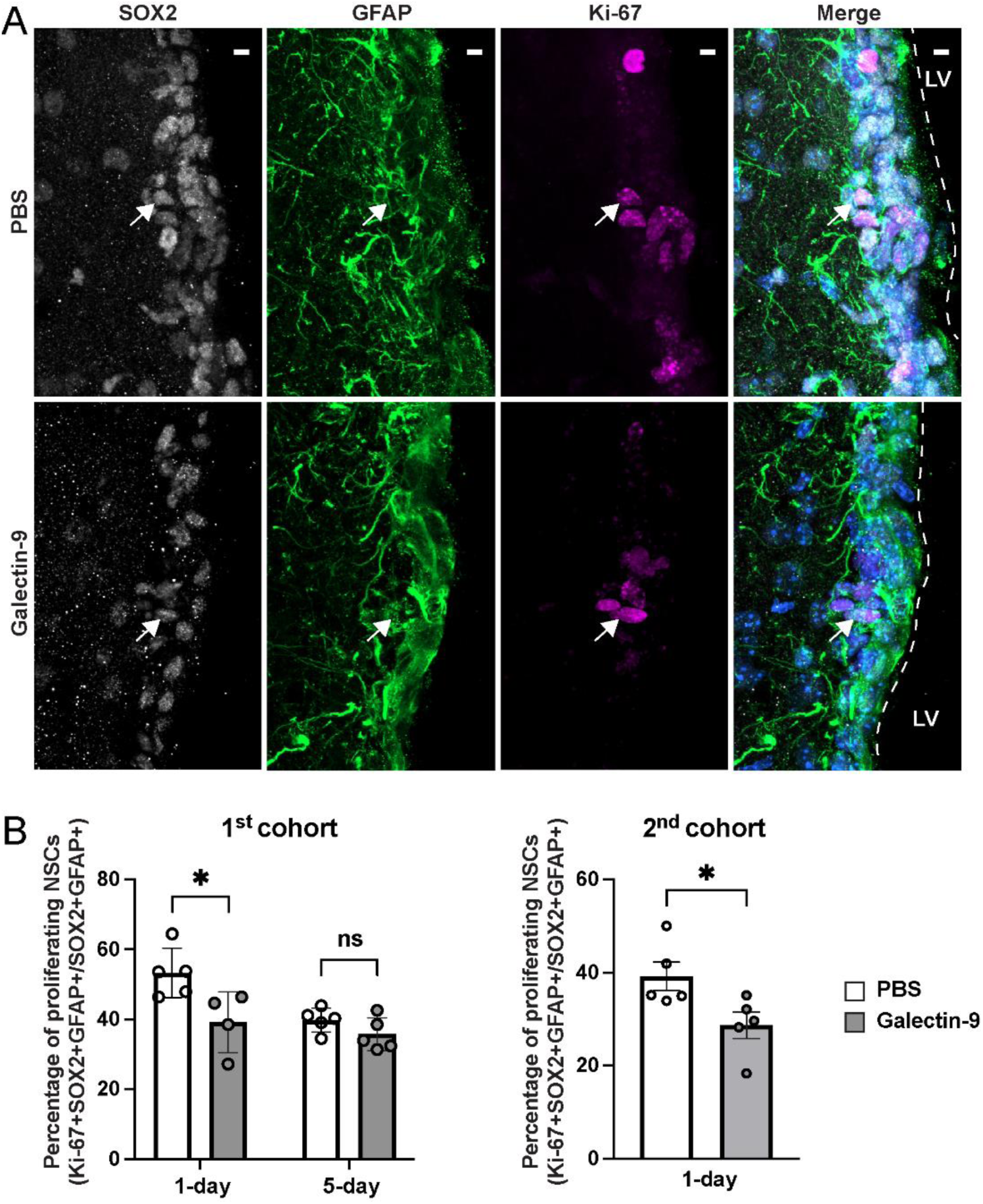
ICV injection of galectin-9 inhibits proliferation of NSCs in the V-SVZ niche of adult mouse brain. (A) Immunostaining of the lateral wall of 8-week-old C57BL/6 mouse brains ICV injected with PBS or galectin-9 for SOX2, GFAP, Ki-67 and merged with Hoechst to label nuclei. SOX2+GFAP+ cells allow identification of NSCs. Ki-67+SOX2+GFAP+ cells represent proliferating NSCs. Arrows indicate representative triple positive cells. Scale bars represent 10 μm. Dotted lines delineate the lateral wall of the lateral ventricle (LV). (B) Quantification of the percentages of proliferating Ki-67+ NSCs in the V-SVZ of 8-week-old C57BL/6 mouse brains ICV injected with PBS or galectin-9 (*N* = 5 for PBS control group in the first cohort, *N* = 4 for the galectin-9 injection group in the first cohort; *N* = 5 per group for the second cohort of experiment). Error bars indicate S.E.M.

### Galectin-9 functions as a negative modulator of the proliferation of NSCs under physiological conditions in the V-SVZ

Based on our findings of the negative impact of galectin-9 on NSC proliferation, we next compared the NSCs in the V-SVZ niche of galectin-9 knockout ^26,46^ to wild type (WT) brains to ask whether galectin-9 controlled NSC proliferation under physiological conditions. The sections of forebrains from 8-week-old *Lgals9*^−/−^ (Gal9-KO) mice and age-matched WT mice were collected and analyzed by immunohistochemistry (Figure 6A and 6B). First, we quantified the proliferating NSCs in the V-SVZ niche. The percentage of proliferating NSCs from Gal9-KO brains was significantly higher compared with those from WT ones (unpaired *t*-test, *t*_(6)_ = 3.792, *P* = 0.009) (Figure 6A and 6C). Moreover, the density of NSCs (marked by SOX2 and GFAP) in the V-SVZ niche of Gal9-KO mice showed a striking increase compared to the WT mice (unpaired *t*-test, *t*_(6)_ = 6.384, *P* = 0.0007) (Figure 6A and 6D). These data strongly suggested that when galectin-9 was absent, NSCs proliferation was enhanced and therefore more NSCs could be produced over the course of time. Thus, galectin-9 might function as a checkpoint molecule, controlling the amount of NSCs in the brain. Compared to the short-term effects of injected recombinant galectin-9, our data presented here evaluated the long-term, chronic role of galectin-9 loss of function on the regulation of NSC proliferation. We further examined the expression of TIM-3, a canonical receptor of galectin-9. The density of the puncta of TIM-3 immunostaining was significantly lower in the Gal9-KO mouse brains compared to that of WT (unpaired *t*-test, *t*_(6)_ = 2.796, *P* = 0.031) (Figure 6B and 6E). Hence, the negative regulatory effects of galectin-9 on NSC proliferation might be mediated by the TIM-3 receptor.

**Figure 6.**
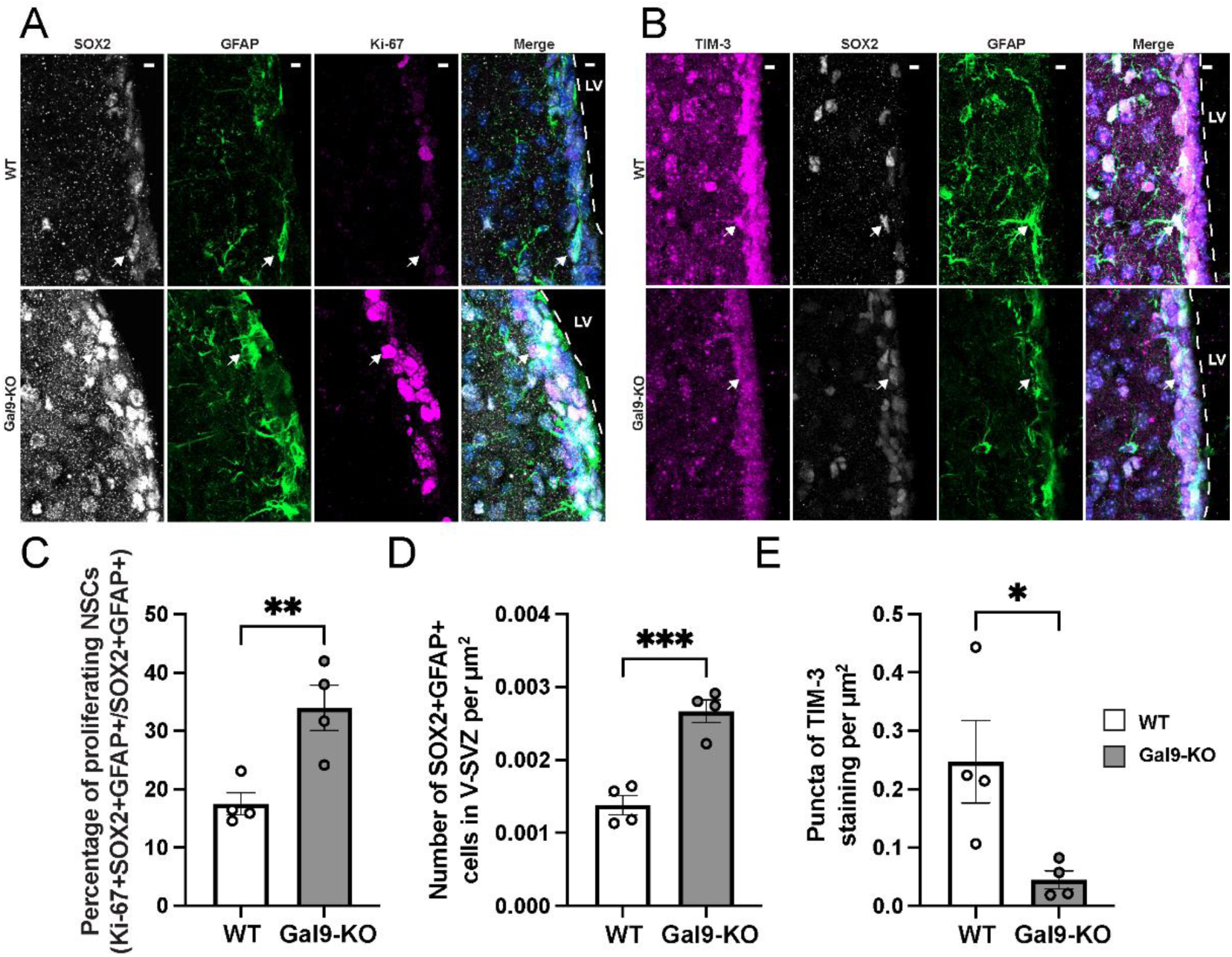
Gal9-KO results in increased proliferation of NSCs in the V-SVZ niche of adult mouse brain. (A) Immunostaining of the lateral wall of age-matched 8-week-old wild type (WT) and Gal9-KO (*Lgals9*^−/−^) C57BL/6 brains for SOX2, GFAP, Ki-67 and merged with Hoechst to label nuclei. SOX2+GFAP+ cells allow identification of NSCs. Ki-67+SOX2+GFAP+ triple positive cells represent proliferating NSCs. Arrows indicate the representative triple positive cell. (B) Immunostaining of the lateral wall of age-matched 8-week-old wild type (WT) and Gal9-KO (*Lgals9*^−/−^) C57BL/6 mouse brains for TIM-3, SOX2, GFAP, and merged with Hoechst to label nuclei. SOX2+GFAP+ cells allow identification of NSCs. Arrows indicate the representative NSC that expresses TIM-3. Scale bars represent 10 μm. Dotted lines delineate the lateral wall of the lateral ventricle (LV). (C) Quantification of the percentages of proliferating Ki-67+ NSCs in the V-SVZ of age-matched 8-week-old wild type (WT) and Gal9-KO (*Lgals9*^−/−^) C57BL/6 mouse brains. (D) Quantification of the densities of SOX2+GFAP+ NSCs in the V-SVZ of age-matched 8-week-old wild type (WT) and Gal9-KO (*Lgals9*^−/−^) C57BL/6 mouse brains. (E) Quantification of the densities of the puncta of TIM-3 staining in the age-matched 8-week-old wild type (WT) and Gal9-KO (*Lgals9*^−/−^) C57BL/6 mouse brain sections. *N* = 4 per group for Figure 6C, 6D and 6E. Error bars indicate S.E.M.

## DISCUSSION

Although a significant body of work has demonstrated enhanced neurogenesis post-ischemic stroke ^12^, signal transduction from the infarct region to the NSC niche has not been systematically investigated. Our study, therefore, set out to examine the intercellular signaling inferred computationally via both *in vitro* and *in vivo* assays. We first computationally inferred intercellular communication based on spatial transcriptomic data. Many of the ligand-receptor pairs identified by CellChat ^22,47,48^ have been validated by previous studies, which affirmed the high quality of our spatial transcriptomic data, and the effectiveness and accuracy of our bioinformatic approach. For instance, one of the ligand-receptor pair we identified was the chemokine SDF-1 (Stromal cell-derived factor 1, also known as CXCL12) and its receptor CXCR4. Previous studies have shown that SDF-1 was expressed in reactive astrocytes and activated microglia after stroke and guided the migration of neural progenitor cells and neuroblasts to the injury site ^12,49^. Another example was *Grn* (or *Pgrn*, Progranulin) that stimulated cell proliferation in the V-SVZ after ischemic stroke and promotes neuronal differentiation in the peri-infarct region ^37,50^. Although we identified a few chemokines that might play important roles in post-ischemic stroke processes, the lack of statistically significant responses to the chemokines from NSCs disassociated from secondary neurospheres in our assay might be due to the NSCs not reaching a migration competent stage and is hence a limitation of this work.

Out of several candidate genes of interest, we for the first time, to our knowledge, reported the evidence of a regulatory role on NSC proliferation by a signaling pathway centered on galectin-9. We showed that galectin-9 inhibited NSC proliferation both *in vitro* (Figure 3) and *in vivo* (Figure 5). We also demonstrated that knocking out galectin-9 resulted in increased NSC proliferation (Figure 6). The molecular mechanism underlying this inhibitory effect has not been characterized to the best of our knowledge. Perhaps contradictory to the concept of mobilization boosting NSCs proliferation post-ischemic stroke to aid neurogenesis ^9^, we identified galectin-9 as a checkpoint molecule which inhibited NSC proliferation. We suggested that galectin-9 might play such a role in the sub-acute and chronic/recovery phase post-ischemic stroke when the proliferation response of NSCs declines.

To investigate the mechanism behind the inhibitory effect of galectin-9 on NSC proliferation, we further examined the potential receptors for galectin-9 identified by our spatial transcriptomics data, including CD44, CD45, TIM-3, IGHM, BCR, and P4HB (Table S1, Figure 1C and 1D). Besides our present work, a few previous studies have identified that galectin-9 plays important roles in the response to stroke injury and explored the receptors involved, including TIM-3 ^27,28^, CD44 ^31^, and TLR4 ^30^. In addition to microglia ^51^, T cell immunoglobulin and mucin-domain containing-3 (TIM-3, encoded by *Havcr2*) has long been known to be expressed in various CNS tissues and upregulated in response to inflammation and infection ^52^. Single-cell analysis showed that TIM-3 expression was found in NSCs, TAPs, and dividing TAPs (Figure 4A). It has been reported that stroke increased the expression of TIM-3 ^27,28^ and galectin-9 ^27–29^. The protective effects of preconditioning ^27^ and postconditioning ^28^ on stroke were mediated by the downregulation of galectin-9/TIM-3 signaling pathway. These data are supportive to the identification of galectin-9 as an upregulated signaling molecules post-ischemic stroke from our spatial transcriptomics data (Figure 1). Moreover, our data demonstrated the downregulation of TIM-3 in Gal9-KO mouse brain (Figure 6B and 6E), which upheld the perspective of galectin-9/TIM-3 signaling pathway participating in the recovery process of stroke injury.

Another receptor identified by our spatial transcriptomics data was CD44 (Table S1). A recent study suggested that galectin-9 produced by microglia promoted remyelination after stroke through CD44 receptor on primitive oligodendrocyte progenitor cells ^31^. However, our results showed that galectin-9 likely originated from ependymal cells (Figure 2B and 4B), which normally express this molecule under physiological conditions. We also demonstrated that galectin-9 suppressed the proliferation of NSCs (Figure 3 and Figure 5) and therefore might potentially negatively impact neurogenesis. One explanation for these seemingly contradictory effects is that our data were collected during a longer time course at different timepoints instead of one single time point (at day-3 post-injury) in the study by Bing Han *et al* ^31^. Although NSC mobilization can be beneficial to the recovery during the time immediately post-stroke, galectin-9 may serve as a break when it is time for the proliferation of NSCs to taper off. Moreover, it is common that one signaling molecule may function in various fashions through divergent pathways at distinct stages of a biological event.

As a immune regulator ^24^, it is natural to postulate the regulatory effects of galectin-9 on microglia, one important type of immune cells in the CNS ^53^. A study showed that galectin-9 may switch its roles from promoting differentiation of microglia into M1-type at the acute phase to directing M2-type differentiation in the later stage through TLR4 receptor to promote recovery post intracerebral hemorrhage in the rat brain ^30^. While TLR4 is highly expressed in microglia populations, our analysis of the existing single-cell dataset showed that the expression of TLR4 demonstrated the second highest level in the cluster of ependymal cells after microglia (Figure 4A). The expression of TLR4 in nonhematopoietic cells of CNS, including ventricular ependymal lining has been experimentally confirmed by *in situ* hybridization previously ^54^ and demonstrated in our current immunohistochemistry data as well (Figure 4D). In addition, the expression of galectin-9 in ependymal cells was also shown in the single-cell analysis (Figure 2B) and experimentally verified by our present study (Figure 4B). It has been shown that the activation of TLR4 by proinflammatory ligand lipopolysaccharide (LPS) could lead to increased galectin-9 expression in human mesenchymal stromal cells (MSCs) ^55^ and human periodontal ligament (PDL)-derived cells ^56^. The galectin-9 synthesized in MSCs in turn inhibited T-cell proliferation ^55^. Mouse brain infection could cause increased expression of TLR4 and TRL2, noticeably in ependymal cells of the choroid plexus ^57^. In the neurocysticercosis mouse brain, TLR4 expression increased in neurons and the expression of TLR2 and TLR7 was upregulated in ependymal cells ^58^. Studies have shown that TLR4 on ependymal cells could respond to systemic inflammatory cytokines ^59^. Not only do Toll-like receptors play important roles in the function of immune cells, many types of cells outside the immune system with immune properties upregulate the expression of TLRs in response to inflammatory stimuli ^60^. Interestingly, the crosstalk between TIM-3 and TLR4 may occur through the PI3K-Akt-mTOR-NF-κB axis ^61^.

Taken together, we propose a working model for the signaling pathway centered on galectin-9 under injured and physiological states (Figure 7). Under physiological conditions, galectin-9 produced by ependymal cells serves to suppress NSC proliferation. Post-ischemic stroke, proinflammatory cytokine(s) are produced at the ischemic stroke infarct site which may bind the TLR4 receptor on ependymal cells and stimulate these cells to increasingly express galectin-9. Galectin-9 can activate the TIM-3 receptor on NSCs and enhance the suppression of NSC proliferation, possibly through mTOR related pathway ^62^ in the later stage of post-ischemic stroke to shut off the initial response of NSCs. Our study pinpoints galectin-9 as a modulator of NSC proliferation based on the landscape of cell-cell communication inference from spatial transcriptomics datasets over a time course of distinctive stages post-ischemic stroke. Further research will focus on the detailed molecular mechanisms underlying galectin-9 signaling and development of the potential therapeutic approaches which leverage this pathway.

**Figure 7.**
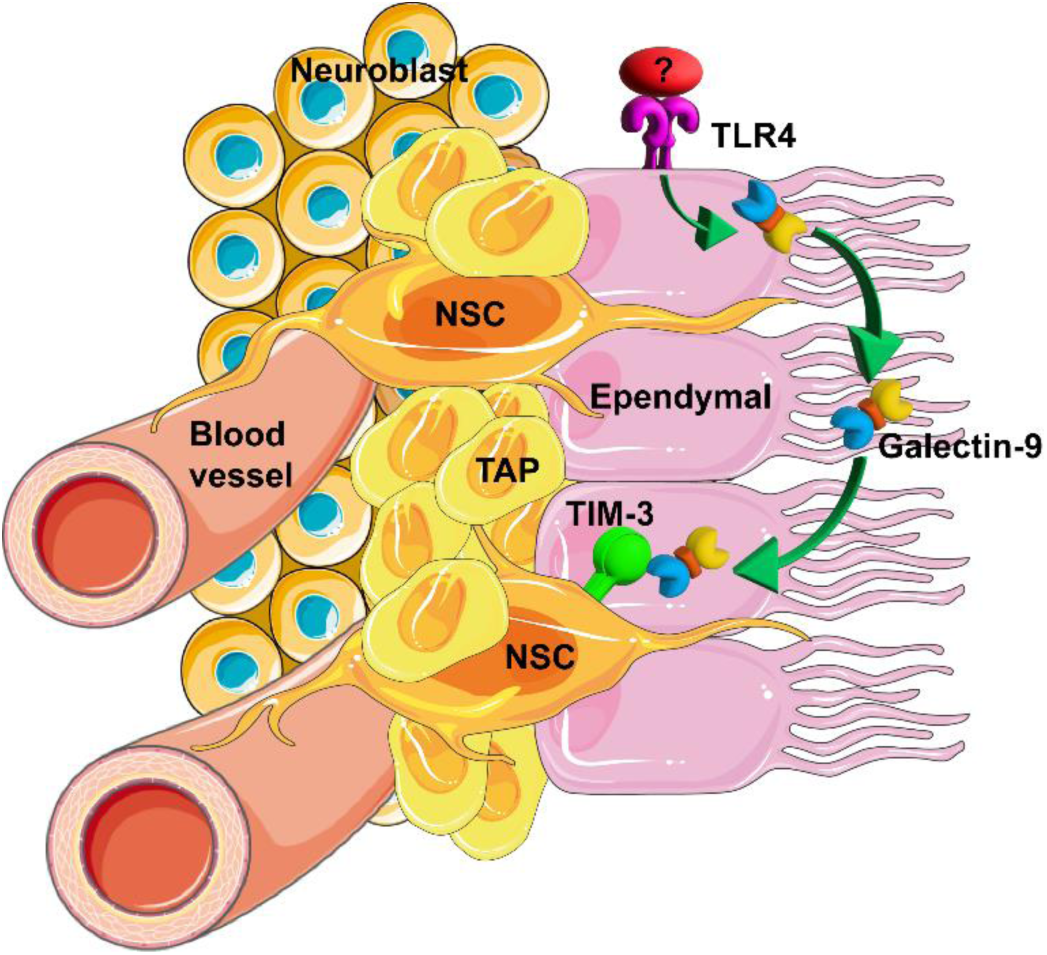
Working hypothesis for the impact of galectin-9 centered signaling pathway on NSCs under physiological conditions and post-ischemic stroke. TLR4 receptors on ependymal cells are activated by proinflammatory cytokines (red oval with a question mark) elicited after stroke. The activated TLR4 signaling pathway triggers ependymal cells, which express galectin-9 under physiological conditions, to increase its expression. Galectin-9 can bind TIM-3 on NSCs and inhibits proliferation to prevent over-proliferation following injury. TAP represents transit-amplifying progenitor; NSC represents neural stem cells.

## METHODS

### Mice

8-week-old male C57BL/6 wild type (WT) mice were obtained from Charles River and housed at the Department of Comparative Medicine at the University of Toronto, following a 12 h light/dark cycle and had *ad libitum* access to water and rodent chow. Age-matched *Lgals9*^−/−^ (Gal9-KO) mice of C57BL/6 genetic background as previously described ^26^, were housed in a specific pathogen-free animal facility at University of Toronto Scarborough, Toronto, Canada. For ICV injections, C57BL6/J mice were ordered from the in-house breeding colony and housed at The Centre for Phenogenomics (Toronto, Canada). All procedures complied with policies formed by the Canadian Council of Animal Care and approved by the local Animal Care Committee at the University of Toronto and The Centre for Phenogenomics, Toronto, Canada.

### Visium spatial transcriptomics

The 10x Visium spatial transcriptomics profiling data were described previously ^63^ and are available at NCBI SRA database BioProject Accession: PRJNA952594) ^63^.

### Inference of cell-cell communication

The Space Ranger output were processed by Seurat (version 5.3.0) ^64^ implemented in R (version 4.5.1) ^65^. The data were then normalized and scaled by SCTransform. Principal components were computed, and clustering was performed with 30 principal components and a resolution of 0.8. UMAP plots were created using the DimPlot and FeaturePlot functions implemented in Seurat (version 5.3.0) ^64^. The stroke infarct regions and the saline injection site of sham control were manually delineated from 10x Genomics Loupe Browser. To identify NSCs, the barcoded spots in the V-SVZ were hand selected from 10x Genomics Loupe Browser. The sum of the normalized counts of the NSC marker genes – *Sox2*, *Nes*, *Gfap*, *Prom1*, and *Fabp7* was calculated and then ranked for all spots. The identity of NSC was assigned to spots in the first quartile of the ranks. Cell-cell communication was inferred by CellChat (version 2.2.0) ^22^ Comparison analysis was conducted among sham, day-2, 10, and 21 post-ischemic stroke for the second dataset following the CellChat analysis pipeline.

### Analysis of scRNA-sequencing data

Whole single-cell RNA sequencing (scRNA-Seq) data of microdissected ventricular-subventricular zone (V-SVZ) on the lateral wall of the lateral ventricle of male (postnatal day 35) and female (postnatal day 29) hGFAP-GFP mice were obtained from NCBI GEO Data (<u>GSE165554</u>). We used the data from the male and female mixed sample (<u>GSM5039270</u>) ^38^. The Seurat object (.rds file) was loaded into R (version 4.5.1) ^65^ and analyzed by Seurat (version 5.3.0)^64^. DimPlot and FeaturePlot implemented in Seurat (version 5.3.0) ^64^ were used to generate the UMAP of clusters of V-SVZ cell types and the expression patterns of selected genes in Figure 2B and 4A.

### Neurosphere cultures

The neuroshpere cultures were prepared as described before ^66,67^. Ventricular-subventricular zone (V-SVZ) on the lateral wall of the lateral ventricle was microdissected from the brains of 8-week-old male C57BL/6 mice. Neurospheres were grown in NeuroCult™ Basal Medium (Mouse & Rat) containing NeuroCult™ Proliferation Supplement (Mouse & Rat) (STEMCELL Technologies), FGF2 (Corning), EGF (STEMCELL Technologies), 0.2% heparin (Sigma-Aldrich, Cat. #: H4784) and 1% penicillin-streptomycin. Following 7-10 days of growth of the primary spheres, spheres were disassociated and seeded for cell proliferation and Boyden chamber assays.

### Cell proliferation assay

The tissue from the V-SVZ of two mouse brains was pooled together as one biological replicate. Four biological replicates for each group were used in one cohort of experiments. The secondary spheres were seeded into each well of a 96-well plate and cultured with 2 μg/ml galectin-9 (R&D Systems, Cat. #: 3535-GA), 200 ng/ml of CCL5 (PeproTech, Cat. #: 250-07), CXCL5 (BioLegend, Cat. #: 573302), and CXCL10 (PeproTech, Cat. #: 250-16), along with control (adding corresponding solvents) for 5 days. Each biological replicate was seeded with technical triplicates in 96-well plates. Cell proliferation was evaluated indirectly using the resazurin reduction assay and quantified by the fluorescence intensity of resorufin ^43^.

### Boyden chamber assay

Cell migration was evaluated by Boyden chamber assay as described previously ^44^. One Millicell^®^ cell culture insert with a pore size of 8 μm (Millipore, Cat. #: PI8P01250) (as the upper chamber) was put into a well of a 24-well cell culture plate (as the lower chamber) to form a Boyden chamber. Both chambers were coated with 40 μg/ml poly-D-lysine (Sigma-Aldrich, Cat. #: P6407-5MG) and 4 μg/ml laminin (Corning, Cat. #: 354232) dissolved in sterile Milli-Q^®^ water. 600 μl coating solution was added to the lower chamber and 400 μl coating solution was added to the upper chamber and then incubated overnight at room temperature. 600 μl of 2 μg/ml galectin-9 (R&D Systems, Cat. #: 3535-GA), 200 ng/ml of CCL5 (PeproTech, Cat. #: 250-07), CXCL5 (BioLegend, Cat. #: 573302), or CXCL10 (PeproTech, Cat. #: 250-16) dissolved in growth factor-free NeuroCult™ Basal Medium (Mouse & Rat) with 10% NeuroCult™ Proliferation Supplement (Mouse & Rat) (STEMCELL Technologies) were placed in the lower chamber, along with controls (media only). 10,000 cells disassociated from the secondary spheres suspended in 400 μl of growth factor-free NeuroCult™ Basal Medium (Mouse & Rat) with 10% NeuroCult™ Proliferation Supplement (Mouse & Rat) (STEMCELL Technologies) were seeded in the upper chamber. The cells dissociated from the secondary spheres cultured from the combination of V-SVZ tissue of two mouse brains were used as one biological replicate for each group in one cohort of experiments. Each biological replicate was seeded with two technical replicates in the Boyden chamber. The chambers were incubated for 6 h at 37°C in a humidified incubator with 5% CO_2_. After the incubation period, the filter was removed, and the upper side of the insert filter, containing non-migrating cells, was removed with a cotton swab, and washed with phosphate buffered saline (PBS, 137 mM NaCl, 2.7 mM KCl, 10 mM Na_2_HPO_4_, 1.8 mM KH_2_PO_4_) three times. The insert filter was then fixed with 4% paraformaldehyde (PFA) in PBS for 20 min, washed with PBS, and stained with Hoechst 33258 (Sigma-Aldrich, Cat. #: B2883) for 10 min protected from light. The membrane of each insert was cut off by a razor blade. The membrane was placed into one drop of PermaFluor™ Aqueous Mounting Medium (Epredia, Cat. #: TA-030-FM) on the uncoated glass slides (Fisherbrand Superfrost Plus Microscope Slides, Cat. #: 1255015), with the side that was facing the migration cues upwards, covered by coverslip, and dried at room temperature in the dark. Cell migration was quantified by counting the migrated cells on each insert filter as described in the “Image Quantification” section below.

### Intracerebroventricular (ICV) injection

Recombinant *Escherichia coli*-derived mouse galectin-9 protein (R&D Systems, Cat. #: 3535-GA) was dissolved in sterile PBS and desalted by Amicon^®^ Ultra Centrifugal Filter, 30 kDa MWCO (MilliporeSigma, Cat. #: UFC503024) at 14,000g. The PBS used for injection into the control mouse brains was centrifuged in parallel with the galectin-9 solution in the same type of filter. The concentration of galectin-9 solution for injection was measured by *DC*™ Protein Assay Kit II (Bio-Rad, Cat. #: 5000112). Prior to stereotaxic surgery, 8-week-old male C57BL/6J mice received a subcutaneous dose of 1.2 mg/kg slow-release buprenorphine for analgesia. Animals were anesthetized with 2-3% isoflurane and received 1 ml of warmed sterile saline subcutaneously in the flank and 8 mg/kg bupivacaine-HCl subcutaneously in the skin overlying the skull prior to being secured in the stereotaxic frame. After removing fur from the scalp and disinfecting the surgical site, a midline incision was made using a #10 scalpel blade and the skin was retracted to expose the skull. A dental drill was used to create a burr hole at 0.5 mm anterior and 1.0 mm lateral to Bregma. 3 µl of galectin-9 (0.1 mg/ml in PBS) was injected into the right lateral ventricle at a depth of 2.3 mm from the skull surface at a rate of 0.1 μl/min using a 10 μl Hamilton syringe fitted with a 29G needle. The control mice received PBS injections. Mouse brains were collected as described in the “Brain tissue preparation and sectioning” section in 1– and 5-day after injection for the first cohort and 1-day post-injection for the second cohort.

### Brain tissue preparation and sectioning

Sections were prepared as described previously ^66^. Mice were anesthetized with 2–3% isoflurane inhalation and then perfused transcardially with PBS followed by 4% PFA. Mouse brains were dissected out, post-fixed for 24-48 h in 4% PFA and then cryoprotected in 30% sucrose in PBS until the brains sank to the bottom of the Falcon™ tube. Tissues were then embedded with Optimal Cutting Temperature (O.C.T.) compound (Fisher Healthcare Tissue-Plus™ O.C.T. Compound, Cat. #: 23730-571). Brain sections were cut 18-μm thick in the coronal plane using a Thermo Fisher Scientific HM525 NX cryostat at −23°C. Sections were collected on glass slides (Fisherbrand™ Superfrost™ Plus Microscope Slides, Cat. #: 12-550-15) coated with gelatin and stored frozen until use.

### Antibodies

The primary antibodies used for immunohistochemistry (IHC) in this study included: goat anti-mouse galectin-9 (R&D Systems, Cat. #: AF3535, RRID: AB_2137240) reconstituted at 0.2 mg/ml in sterile PBS and diluted at 1:30; mouse anti-FOXJ1 (eBioscience™, Cat. #: 14-9965-80, RRID: AB_1548836) at 1:200 dilution; biotin anti-mouse CD366 (Tim-3) (BioLegend, Cat. #: 119720, RRID: AB_2571936) at 1:150 dilution; rabbit anti-TLR4 (Proteintech, Cat. #: 19811-1-AP, RRID: AB_10638446) at 1:200 dilution; rabbit anti-SOX2 (Cell Signalling Technology, Cat. #: 3728S, RRID: AB_2194037) at 1:250 dilution; goat anti-SOX2 (R&D Systems, Cat. #: AF2018, RRID: AB_355110) reconstituted at 1 mg/ml in sterile PBS and diluted at 1:100; mouse anti-GFAP (Cell Signalling Technology, Cat. #: 3670S, RRID: AB_561049) at 1:500 dilution; chicken anti-GFAP (MilliporeSigma, Sigma-Aldrich, Cat. #: AB5541, RRID: AB_177521) at 1:500 dilution; mouse anti-Ki-67 (BD Biosciences, Cat. #: 550609, RRID: AB_393778) at 1:100 dilution.

The secondary antibodies used for IHC in this study included: Alexa Fluor^®^ 488-conjugated donkey anti-chicken secondary antibody (Jackson ImmunoResearch Laboratories, Cat. #: 703-545-155, RRID: AB_2340375) at 1:500 dilution; Alexa Fluor^®^ 647-conjugated donkey anti-rabbit secondary antibody (Jackson ImmunoResearch Laboratories, Cat. #: 711-605-152, RRID: RRID: AB_2492288) at 1:500 dilution; Cy™3-conjugated donkey anti-mouse secondary antibody (Jackson ImmunoResearch Laboratories, Cat. #: 715-165-150, RRID: AB_2340813) at 1:500 dilution; Alexa Fluor^®^ 488-conjugated donkey anti-goat secondary antibody (Jackson ImmunoResearch Laboratories, Cat. #: 705-545-147, RRID: AB_2336933) at 1:500 dilution; Cy™3 Streptavidin (Jackson ImmunoResearch Laboratories, Cat. #: 016-160-084, RRID: AB_2337244) at 1:1000 dilution.

### Immunohistochemistry (IHC)

Frozen tissue sections were dried for 30 min at 37°C and then rehydrated. Sections were then washed three times with washing buffer (0.1% Triton X-100 in PBS) for 5 min each time. Subsequently, sections were blocked and permeabilized in the blocking solution of 3% donkey serum (Jackson ImmunoResearch Laboratories, Cat. #: 017-000-121, RRID: AB_2337258, reconstituted in Milli-Q^®^ water at 60.0 mg/ml) and 0.3% Triton X-100 in PBS for 1 h. Then sections were incubated in primary antibodies listed in the “Antibodies” section diluted in the blocking solution overnight at 4°C in a humidified chamber. The following day, sections were washed three times in washing buffer for 5 min per wash, then incubated in corresponding secondary antibodies listed in the “Antibodies” section diluted in PBS for 2 h at room temperature in the dark. Afterwards, sections were washed three times in the washing buffer for 5 min per wash. Finally, sections were counterstained for 5 min at room temperature in 0.5 μg/ml Hoechst 33258 (Sigma-Aldrich, Cat. #: B2883), washed with PBS for 5 min, and then mounted with PermaFluor™ Aqueous Mounting Medium (Epredia, Cat. #: TA-030-FM) and dried overnight at room temperature.

### Image acquisition

Neurosphere images were obtained using a Zeiss Apotome live cell system with Axiocam 506 monochrome camera and 10 × objective at the University of Toronto Temerty Faculty of Medicine Microscope Imaging Laboratory. Each entire well of the 96-well plate was imaged for subsequent quantification. Immunofluorescence images were collected using a Zeiss Axio Observer 7 equipped with LSM 800 scan head with 60 × objective at the Collaborative Advanced Microscopy Laboratories of Dentistry (CAMiLoD) at the University of Toronto Faculty of Dentistry. Images were acquired using Z-stacks at 1 μm interval in Zen Blue (version 2.6). All images were acquired with Z-stack sizes ranging between 17-22 slices. Images shown were produced using the orthogonal projection feature implemented in Zen Blue.

### Image quantification

Image quantifications were all done using Fiji (ImageJ2, version 2.16.0/1.54p) ^68^. The diameter of the neurospheres (Figure 3C) was quantified by “Measure” function implemented in Fiji and then converted to μm according to the scale bar obtained in Zen Blue. The number of neurospheres (Figure 3D) was quantified by manual counting for each well of a 96-well plate. Neurospheres cultured from the combined tissue from the V-SVZ of two mouse brains was pooled together as one biological replicate. Four biological replicates were quantified for each group in one cohort of experiment and technical triplicates were quantified for each biological replicate as described in the “Cell proliferation assay” section.

For quantification of the images from the Boyden chamber assay (Figure S1), the nuclei staining images of the insert filter were obtained under 10 × objective. A region of interest (ROI) with an identical size of 3360 × 3468 pixels covering most part in the center of the image was selected for all images in Fiji. The number of pixels in the ROI were calculated and converted to mm^2^ according to the scale bar obtained in Zen Blue. All migrated cells represented by the stained nuclei inside the ROI were counted. The migrated cells were presented by dividing the numbers of cells in the ROI of the insert filter on the side facing the migration cues by the area of the ROI. The cells dissociated from the secondary spheres cultured from the combination of V-SVZ tissue of two mouse brains were used as one biological replicate for each group in one cohort of experiment. Each biological replicate was seeded with two technical replicates in the Boyden chamber.

For quantification of the proliferating NSCs in the V-SVZ (Figure 5B and 6C) only visibly immunostained cells along the dorsal and ventral portions of the lateral wall (LW), closest to the ventricle (periventricular area) were counted. The anatomically matched sections were quantified for the groups of control and treatment. To obtain a cell count value for each individual brain as one biological replicate, the positive cells counted from five anatomically matched 18-μm thin coronal sections containing the V-SVZ LW region were totaled and averaged. The percentage of proliferating (Ki67+) NSCs was represented by dividing the number of SOX2+GFAP+Ki-67+ cells by the number of SOX2+GFAP+ cells.

For quantification of the density of the NSCs in the V-SVZ (Figure 6D), the V-SVZ at the LW was manually selected in Fiji. The number of pixels in the selected area were quantified by Fiji and converted to μm^2^ according to the scale bar obtained in Zen Blue. Cells inside the selected V-SVZ area were counted. SOX2+GFAP+ cells represent NSCs. Then the density of NSCs in the V-SVZ was calculated by dividing the number of NSCs by the area of the selected V-SVZ region. To obtain a density of cell value for each individual brain as one biological replicate, the positive cells counted from three anatomically matched 18-μm thin coronal sections containing the V-SVZ LW region were totaled and averaged.

The quantification of the density of the puncta of TIM-3 staining (Figure 6E) was done by automatic particle counting in Fiji. A ROI with a size of 1500 × 2000 pixels covering most part of the image field obtained by a Zeiss Axio Observer 7 equipped with LSM 800 scan head with 60 × objective was selected for analysis for all images of anatomically matched 18-μm thin coronal sections containing the V-SVZ LW region. The number of pixels in the ROI were calculated and converted to μm^2^ according to the scale bar obtained in Zen Blue. All images were processed by an identical process written in a Macros implemented in Fiji of steps as follows. The image was converted to 8-bit. The thresholds were set to (140, 255). The number of puncta was counted by “Analyze Particles” function implemented in Fiji. The density of the puncta of TIM-3 staining was calculated by dividing the number of puncta by the area of the ROI. To obtain a density of puncta value for each individual brain as one biological replicate, three anatomically matched 18-μm thin coronal sections containing the V-SVZ LW region were analyzed, totaled and averaged.

### Statistics

All statistical analyses and graphical representations were done in GraphPad Prism software (version 10.5.0 for macOS). For Figure 3B, a Friedman test followed by Dunn’s multiple comparisons test were conducted. For Figure 3C and 3D, a paired *t*-test was used. For Figure 5B, 6C, 6D, and 6E, the unpaired *t*-test was used. For Figure S1, a Friedman test was conducted.

## SUPPLEMENTAL INFORMATION

**Figure S1.**
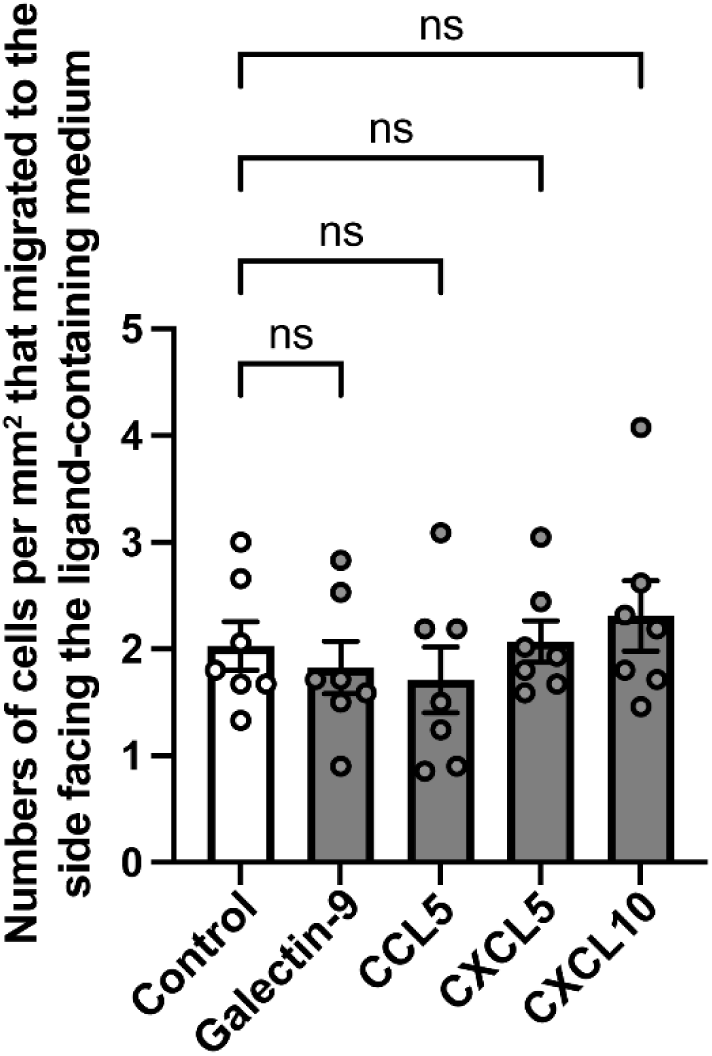
Evaluation of the select signaling molecules on NSC migration. The numbers of cells per mm^2^ that migrated to the side of the Boyden chamber facing the ligand-containing medium were quantified. *N* = 7 per group. Error bars indicate S.E.M.

**Table S1. Pairwise comparisons of up– and down-regulated signaling ligand-receptor pairs by CellChat inference from spatial transcriptomics data of sham, day-2, –10 and –21 post-ischemic stroke brain sections.**

## RESOURCE AVAILABILITY

### Lead contact

Further information and requests should be directed to the lead contact, Scott A. Yuzwa.

### Materials availability

All materials generated in this study are available from the lead contact.

### Data and code availability

The sequencing data have been deposited in the NCBI SRA database under BioProject Accession: <u>PRJNA952594</u>

## ACKNOWLEDGEMENTS

This work was supported by Canadian Institutes of Health Research (CIHR; grant number PJT-175137 to S.A.Y.). Additional operating funding was provided by a Temerty Pathway grant and a Connaught New Investigator Award to S.A.Y. Infrastructure to support these studies was funded by the Canadian Foundation for Innovation (CFI) and the Ontario Research Fund (ORF). H.H. was supported by University of Toronto Fellowship and Mary H. Beatty Fellowship at the University of Toronto. We thank the Arturo Alvarez-Buylla laboratory for making the R objects publicly available to allow re-analysis of V-SVZ scRNA-seq data. We acknowledge the Collaborative Advanced Microscopy Laboratories of Dentistry (CAMiLoD) and the Faculty of Dentistry, University of Toronto, Toronto, ON, Canada for service, training and expert advice received with microscopy imaging.

## AUTHOR CONTRIBUTIONS

S.A.Y. and H.H. conceptualized and designed the experiments. H.H. performed the neurosphere, cell proliferation, Boyden chamber, tissue sectioning, immunohistochemistry experiments, and image acquisition and quantification. E.D. performed the intracerebroventricular injection. H.H., W.C.J.L., and T.T. analyzed the 10x Visium spatial transcriptomics profiling data. H.H. generated visualizations. D.L.C. and M.F. provided the 10x Visium spatial transcriptomics profiling data. H.C. and B.T. provided the Gal9-KO mouse brains. H.H. and S.A.Y. wrote the initial draft of the paper. All authors commented on and approved the final version of the manuscript before submission.

